# Predicting Prognostic Bidirectional Molecular Signatures Associating Myocardial Infarction and Lung Cancer: An *In-Silico* Perspective

**DOI:** 10.1101/2024.11.15.623806

**Authors:** Dhruva Nandi, Rajiv Janardhanan, Piyush Agrawal

## Abstract

Myocardial Infarction (MI) and lung cancers substantially contributors to morbidity and mortality. They share common risk factors such as smoking, and hypertension. There is a pressing need to identify bidirectional molecular signatures linking MI and lung cancer relationship to create a unique triage to improve the clinical outcomes of the patient. In the current work, we extracted the common differentially expressed genes (DEGs) between MI and lung cancer and identified 2339 upregulated and 2973 downregulated genes in MI datasets; 952 upregulated and 653 downregulated genes in LUAD; 1466 upregulated and 2816 downregulated genes in LUSC. Among these, 27 genes (11 upregulated and 16 downregulated) were common across MI, LUAD, and LUSC. Functional enrichment analysis revealed shared biological processes, such as inflammatory response, and cell differentiation. KEGG pathway analysis highlighted common pathways such as B cell receptor signaling, TNF signaling. A protein-protein interaction study with STRING revealed a variety of interaction partners; miRNA-mRNA network analysis characterizes conserved binding sites for 6 genes while overall survival analysis for lung cancer patients shows a significant association with prognosis. In addition, machine learning models built using the Support Vector Machine and Random Forest algorithms demonstrated high AUROC values of 0.79 and 0.81 respectively on balanced dataset in the classifying MI patients from non-MI patients. Lastly, based on drug repurposing analysis, we proposed FDA-approved drugs such as Venetoclax, Lomitapide, Regorafenib that could potentially target these genes, indicating novel therapeutic options for the co-occurring conditions of MI and lung cancer. Our findings highlight the similarities in molecular makeup between lung cancer and MI, providing information for future investigations and therapeutic approaches.

## Introduction

The most severe clinical manifestation of coronary artery disease (CAD) and one of the most dangerous coronary events associated with sickle cell disease (SCD) is myocardial infarction (MI) [1,2]. The two types of this pathophysiology are non-ST-elevation MI (NSTE-MI) and ST-elevation MI (STE-MI) [3]. Every year, over 3 million people have STE-MI, and over 4 million people have STE-MI pathology. While MI is mostly seen in developed countries, it has been found in developing countries as well [4–7]. Lung cancer is the second most frequently diagnosed malignancy worldwide [8]. It is estimated that over 2.2 million new lung cancer cases and 1.7 million lung cancer-associated fatalities occur annually worldwide [9]. Lung carcinoma is classified into two primary types: small-cell lung cancer (SCLC) and non-small-cell lung cancer (NSCLC). The latter is further separated into three primary histological subtypes: large cell carcinoma (LCC), squamous cell carcinoma (SCC), and adenocarcinoma (ADC) [10].

Globally, acute myocardial infarction (AMI) and cancers are substantial contributors to morbidity and mortality [11]. Research has shown that patients diagnosed with cancer are at a higher short-term risk of experiencing cardiovascular (CV) events, while those with acute myocardial infarction have an increased incidence of cancer, despite the limited available references on the relationship between these two conditions [12,13]. These prospective connections suggest that a latent connection exists between the incidence of AMI and cancer survival. Consequently, it is of the utmost significance to identify common biomarkers for the prediction of AMI and cancer survival. Research has demonstrated that cardiac complications, including myocardial infarction, heart failure, and arrhythmias, are more prevalent among cancer survivors than in the general population [14]. For example, the development of adverse cardiac events in cancer patients may be influenced by inflammation, oxidative stress, and endothelial dysfunction, which are prevalent processes in both cancer and cardiovascular diseases [15].

Many risk factors are shared between cardiovascular disease (CVD) and lung cancer, including smoking, hypertension, diabetes mellitus (DM), advanced age, obesity, and racial and socioeconomic status (SES) [16–18]. Patients with lung cancer frequently have pre-existing CVD’s comorbidities as a result of these overlapping risk factors. In patients with lung and bronchus cancer, hypertension, arrhythmia, CAD, dyslipidemia, and heart failure (HF) were identified as the most prevalent CV conditions [19]. Also, the overall survival rate has been documented as lowest among patients with NSCLC and comorbid coronary heart disease, MI, or cardiac arrhythmias [20]. Lung cancer has been identified as an independent risk factor for the development of CVDs, specifically CAD and MI. Hence, this study aims to identify bidirectional molecular signatures linking MI and lung cancer relationship to create a unique triage to improve the clinical outcomes of the patient. Understanding these shared pathways can offer insights into overlapping disease mechanisms, improving early diagnosis and targeted therapies. Moreover, integrating these signatures may enhance personalized medicine approaches for individuals at risk of both diseases.

## Methodology

### A. Dataset Creation and Processing

We downloaded two transcriptomic datasets, GSE59867 [21] and GSE62646 [22], of MI patients from the GEO database. In the first dataset, i.e., GSE59867, there were 111 MI patients with ST-segment elevation myocardial infarction (SEMI) and 46 patients with stable coronary artery disease (CAD) as the control group. In the second dataset, GSE62646, there were 28 patients with SEMI and 14 patients as part of the control group with stable CAD. For lung cancer, we downloaded RNASeq data for both lung cancer types, LUAD [23] and LUSC [24], profiled in TCGA. HTSeq-count and the ID/gene mapping files were downloaded from the “DATA SETS” page of the UCSC Xena Browser [25] database for both cancer types. In the case of LUAD, we obtained 527 patients and 59 normal samples, whereas in the case of the case of LUSC, we obtained 501 patients and 49 normal samples. As there were multiple transcripts of the same gene in the datasets, we took the average read count for those genes before doing differential analysis.

### B. Characterization of DEGs

We computed the DEGs using the ‘limma’ package [26], which runs on the default parameters. Genes with the FDR of <0.05 were further used for analysis. In the case of MI data, LogFC ranges from 2 to −2; hence, we used the cutoff value ‘0.2’, i.e., genes with a LogFC value >=0.2 or above were termed ‘upregulated’ and genes with a LogFC value<=−0.2 or below were called ‘downregulated’. We chose the cutoff values of LogFC >=2 and −2 to label genes as upregulated and downregulated, respectively, in the TCGA-LUAD and TCGA-LUSC cases, where LogFC ranges from 10 to −10.

### C. Gene Ontology Analysis

We used the Enrichr website (https://maayanlab.cloud/Enrichr/) [27] to characterize enriched biological processes and molecular functions. We used our list of genes (common DEGs obtained in MI, LUAD, and LUSC) as the foreground, and the webserver’s default geneset as the background. We used default parameters for performing the analysis and further considered processes for analysis that were statistically significant (FDR <= 0.1). We also identified the enriched KEGG pathways using the Enrichr tool, where the current version of the KEGG database used is 2021 and is for humans. We used the ‘ggplot2’ R package to display the top 20 enriched pathways as a pictorial representation.

### D. Protein-Protein Interaction Analysis

We performed protein-protein interaction analysis and characterized interacting partners using the STRING database [28], accessible at https://string-db.org/. We provided the gene list as an input and selected ‘Homo sapiens’ as the organism to search for interacting partners. We ran the tool with default parameters, selecting ‘full STRING network’ as the network type, where the edges indicate both functional and physical protein associations, and ‘high confidence (0.70)’ as the minimum interaction required score. We downloaded the output network images in the high-resolution ‘png’ format.

### E. Survival Analysis

To perform survival analysis, we used the GEPIA2 webserver (http://gepia2.cancer-pku.cn/#survival) [29] which performs survival analyses based on gene expression level. We performed the analysis using both individual gene expression as a feature and all genes as gene signatures. The server used the log-rank test, also known as the Mantel-Cox test, for the hypothesis evaluation. Here, we conducted an ‘Overall Survival’ analysis using the ‘Median’ as the group cutoff, classifying 50% of the samples as ‘High’ and the remaining 50% as ‘Low’. We computed the hazard ratio using a 95% confidence interval with ‘Months’ as the axis unit. We considered the genes to be significantly associated with survival if the log rank p-value was <=0.05. We conducted survival analysis for both LUAD and LUSC.

### F. miRNA-mRNA relationship analysis

Understanding miRNA-mRNA interactions is crucial from various perspectives, such as understanding gene regulation, designing miRNA-based therapeutics, evolutionary analysis, etc. Here, we used TargetScanHuman 8.0 (https://www.targetscan.org/vert_80/) [30] software to identify potential miRNAs against the target of interest. The results provided a list of miRNAs and their conserved sites, which match their seed regions. We analyzed the most prevalent transcript results in cases where more than one transcript was present. We ran the analysis using the default settings. The analysis yielded a table containing a list of miRNAs with conserved binding sites on the mRNA, along with the position information of gene 3’UTR and the site type (8 mer, 7 mer, or 6 mer). The table also includes the Context++ score, the weighted context++ score, and the predicted relative K_D_.

### G. Machine Learning Analysis

We developed support vector machine (SVM) models to classify MI patients from the normal population using gene expression as a feature. We performed machine learning analysis using the Python-based ‘Scikit’ package [31]. We used the five-fold cross-validation technique for training and testing and validated the model’s performance on an independent dataset. We performed model validation using both an imbalanced (actual positive and negative cases) and balanced (similar positive and negative cases) dataset. To obtain the best performance, we optimize the parameters ‘kernels’, ‘g’, and ‘C’ of the model during training and testing. We computed the performance in terms of Area Under Receiver Operating Characteristics (AUROC) and created an AUC plot using the R package ‘pROC [32].

### H. Drug Repurposing Studies

We performed a drug repurposing study to propose novel drugs against our choice of targets. Firstly, we downloaded the 3D structure of the targets from the RCSB-PDB database [33]. Next, using the ‘Open Babel’ software [34], we prepared the target for docking. We uploaded the prepared 3D structure of the target to the webserver ‘DrugRep’ [35] for virtual screening and docking analysis. We selected the “protein box size” and the “centers of X, Y, and Z coordinates” after uploading the structure. Finally, we chose the library of ‘FDA-approved drugs’ for the virtual screening and downloaded the results as a zip file containing docked protein-ligand structures and the free energy.

## Results

### 3.1 MI and Lung cancer shares common differentially expressed genes

To characterize the DEGs in the MI and Lung cancer datasets, we used the R package ‘limma’. **Table 1** shows the complete statistics of the results. We used the R packages ‘ggplot2’ and ‘ggrepel’ to visualize the differential gene expression results in the form of a volcano plot. **Fig. 1A&B** shows the result for the 2 MI datasets; **Fig. 1C** shows the result for the TCGA-LUAD dataset; and **Fig. 1D** shows the result shows the result for the TCGA-LUSC dataset. The top 10 differentially up- and down-regulated genes based on LogFC values have been plotted as well in each plot. Based on LogFC cutoff value ‘0.2’ for MI datasets, we obtained 469 up and 895 downregulated genes in GSE59867 and 2210 up and 2707 downregulated genes in GSE62646. We further merged these results from both datasets and obtained unique 2339 upregulated and 2973 downregulated genes. Similarly, for TCGA-LUAD and TCGA-LUSC, a LogFC cutoff of ‘2’ was used to get DEGs, which resulted in 952 up and 653 downregulated genes in LUAD and 1466 up and 2816 downregulated genes in LUSC. The complete results of differential expression analysis for all 4 datasets (2 MI, 1 TCGA-LUAD, and 1 TCGA-LUSC) are provided in **Supplementary Tables S1–S4.**

**Figure 1.**
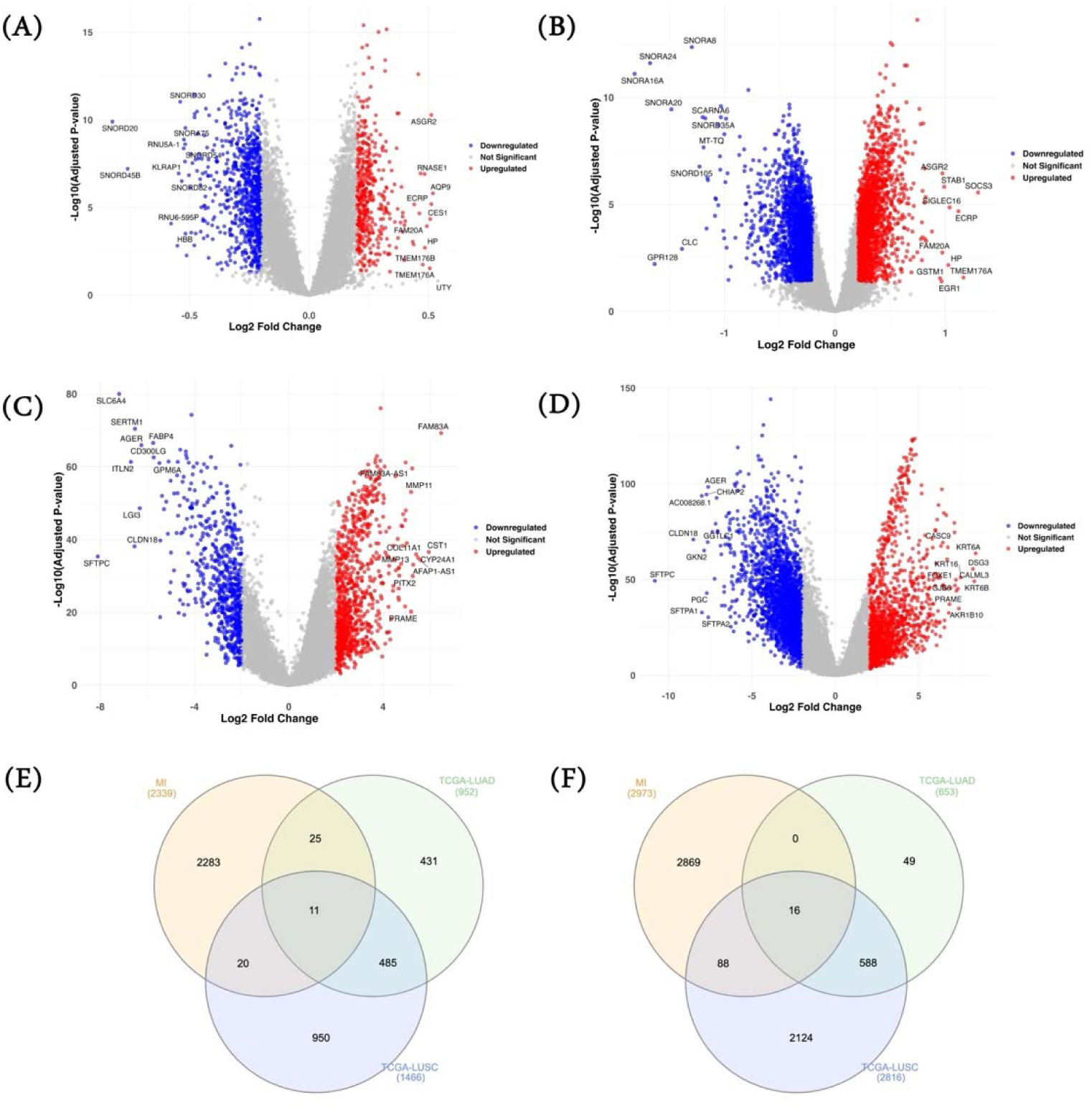
DEGs and common gene characterization. DESeq2 was used to characterize differentially expressed genes on MI datasets (A) GSE and (B); (C) TCGA-LUAD and (D) TCGA-LUSC. Venn diagram showing common upregulated genes among MI datasets (combined), TCGA-LUAD and TCGA-LUSC (E); and common downregulated genes among MI datasets (combined), TCGA-LUAD and TCGA-LUSC (F).

**Table 1.** Differentially Expressed Genes Analysis. ‘limma’ was implemented to compute the differentially expressed genes in MI and Lung cancer datasets.

| Sr. No. | Condition | Dataset Source | Log2FC<br>_Cut off | padj_<br>Cutoff | Upregulate<br>d Genes | Downregulate<br>d Genes |
| --- | --- | --- | --- | --- | --- | --- |
| 1 | MI | GSE59867 | 0.2 | 0.05 | 469 | 895 |
| 2 | MI | GSE62646 | 0.2 | 0.05 | 2210 | 2707 |
| 3 | LUAD | TCGA | 2 | 0.05 | 952 | 653 |
| 4 | LUSC | TCGA | 2 | 0.05 | 1466 | 2816 |

\* MI: Myocardial Infarction; LUAD: Lung Adenocarcinoma; LUSC: Lung Squamous Cell Carcinoma; TCGA: The Cancer Genome Atlas

Next, we overlapped the three sets of differentially expressed genes to obtain the common up- and down-regulated genes **[Figs. 1E and F]**. In total, we obtained 27 common genes (11 upregulated genes and 16 downregulated genes) among MI, LUAD, and LUSC. A list of these common genes is provided in **Table 2**.

**Table 2.** List of Common Up and Downregulated Differentially Expressed Genes among MI, TCGA-LUAD and TCGA-LUSC.

| Sr No. | Up regulated | Down Regulated |
| --- | --- | --- |
| 1 | ASF1B | ACKR4 |
| 2 | B3GNT3 | ALAS2 |
| 3 | CST1 | CA1 |
| 4 | E2F2 | FGFBP2 |
| 5 | EFNA3 | GRIK4 |
| 6 | FGF11 | HBA2 |
| 7 | HMGA1 | HBB |
| 8 | MIOX | IL1RL1 |
| 9 | PLK1 | IL5RA |
| 10 | TERT | LRRK2 |
| 11 | ZP3 | MS4A2 |
| 12 |  | RGS9 |
| 13 |  | SLC14A1 |
| 14 |  | SLC4A1 |
| 15 |  | ST8SIA6 |
| 16 |  | TGFBR3 |

\* MI: Myocardial Infarction; LUAD: Lung Adenocarcinoma; LUSC: Lung Squamous Cell Carcinoma; TCGA: The Cancer Genome Atlas

### 3.2 Functional Enrichment Analysis elucidates common pathways between MI and Lung Cancer

We performed the functional enrichment analysis using the DEGs characterized in each diseased condition and the DEGs common to all three conditions. Processes such as “Positive Regulation of DNA-binding Transcription Factor Activity,” “Regulation of Inflammatory Response,” and “Positive Regulation of NIK/NF-kappaB Signaling” primarily enriched for the upregulated DEGs in MI. **[Fig 2A]**. DEGs that were upregulated in LUAD were linked to processes such as “Positive Regulation of MAPK Cascade”, “Positive Regulation of Protein Phosphorylation”, “Regulation of Angiogenesis”, “Inflammatory Response” and more. **[Fig 2B].** Similarly, in LUSC, processes such as “Epidermis Development”, “Mitotic Spindle Assembly Checkpoint Signaling” and “Epidermal Cell Differentiation” were associated with the upregulated genes. **[Fig 2C]**. **Figure 2(A-C)** lists the top 20 enriched biological pathways, while **Supplementary Tables S5–S7** provide a complete list of enriched biological processes. KEGG analysis reveals processes such as “osteoclast differentiation,” “B cell receptor signaling pathway,” “TNF signaling pathway,” “small cell lung cancer,” etc. in MI. **[Fig 2D]**. In the case of LUAD, we observed pathways like the “cell cycle”, the “p53 signaling pathway”, the “Fanconi anemia pathway”, the “ECM-receptor interaction pathway”. etc. **[Fig 2E]**. In the case of LUSC, only one KEGG pathway was significant, i.e., “cell cycle.” **Supplementary Tables S8–S10** provide a complete list of KEGG pathways. We also performed functional enrichment analysis for downregulated genes, and **Supplementary Figure S1(A-F)** and **Supplementary Table S11-S16** provide all the results.

**Figure 2.**
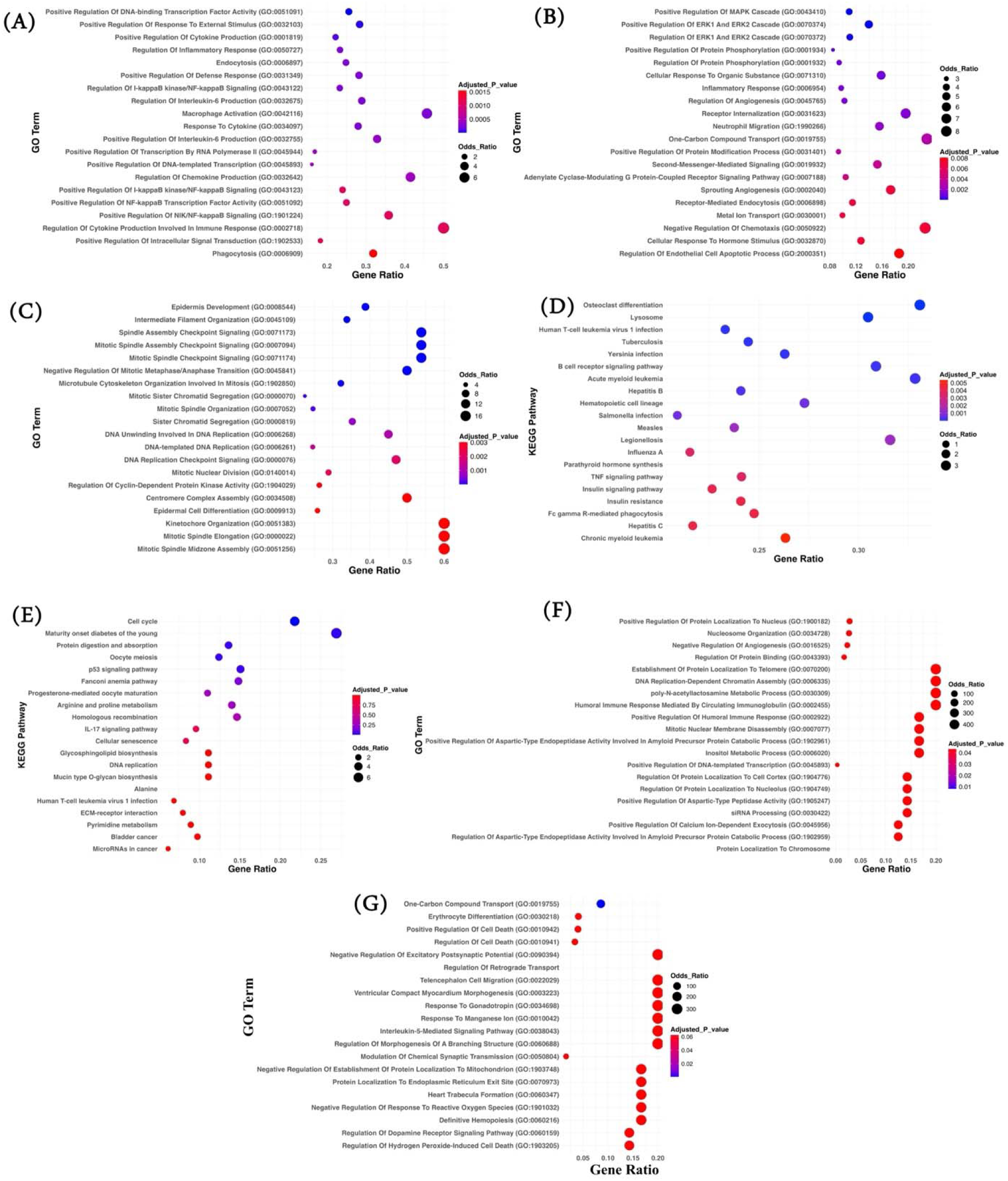
Gene Enrichment Analysis. Panel A-C represents top20 statistically significant enriched biological processes associated with (A) Upregulated genes in MI datasets (combined); (B) Upregulated genes in TCGA-LUAD; (C) Upregulated genes in TCGA-LUSC. Panel D-G represents top20 statistically significant enriched KEGG pathways associated with (D) Upregulated genes in MI datasets (combined); (E) Upregulated genes in TCGA-LUAD; (F) Common 11 upregulated genes among MI, TCGA-LUAD and TCGA-LUSC datasets; (G) Common 16 downregulated genes among MI, TCGA-LUAD and TCGA-LUSC datasets.

Next, we looked for the enriched biological pathways associated with the genes common among the three conditions. Pathways enriched for the common 11 upregulated genes include “Positive Regulation of Protein Localization to the Nucleus”, “Negative Regulation of Angiogenesis”, “Positive Regulation of Humoral Immune Response”, etc. **[Fig. 2F]**. Independent studies have demonstrated the role of protein localization in responding to stress and damage in MI and lung cancer patients. In MI, the heart undergoes significant stress and damage, leading to nuclear signaling pathway activation, which is essential for cell survival and repair [36]. Similarly, in the case of lung cancer, aberrant nuclear localization of proteins can lead to cancer progression, metastasis, and therapy resistance [37]. **Supplementary Table S17** provides a comprehensive list of enriched biological processes. The KEGG analysis revealed enrichment in pathways such as “cell cycle”, “non-small cell lung cancer” and so on. **Supplementary Table S18** provides a complete list of enriched KEGG pathways.

Likewise, for the common 16 downregulated genes, we observed enrichment of processes such as “erythrocyte differentiation”, “regulation of cell death”, “interleukin-5-mediated signaling pathway”, etc. **[Fig. 2G].** The erythrocyte differentiation process has been observed in both conditions previously. In MI patients, the heart undergoes severe damage and stress, and the production of inflammatory cytokines such as TNF-α, IL-1β and IL-6 takes place. These cytokines inhibit the process of erythropoiesis by remodeling the bone marrow microenvironment [38]. In the case of lung cancer, a hypoxic environment is a well-observed phenomenon that alters the normal hematopoiesis as cancer cells compete for oxygen, leading to a less favorable environment for erythrocyte formation and differentiation [39]. A complete list of enriched biological processes is provided in **Supplementary Table S19**. KEGG analysis revealed enrichment of pathways such as “cytokine-cytokine receptor interaction”, “nitrogen metabolism,” etc. A complete list of enriched KEGG pathways is provided in **Supplementary Table S20.**

### 3.3 Protein-Protein Interaction (PPI) Analysis elucidates diverse interacting partner of key targets

We conducted a PPI analysis of common DEGs using the STRING database to characterize their interactions. First, we performed the analysis using the 11 common upregulated DEGs and found that these genes do not form any interaction network in general and are independent. The only interaction observed was among the genes *PLK1* and *ASF1B* **[Fig. 3A]**. The above observation explains that this gene set is heterogeneous, and genes are involved in different biological processes and pathways. Because of this, we looked at each gene separately and saw that they were interacting with many other genes that might be controlling different pathways **[Fig. 3B–L]**. We also conducted a similar analysis using 16 common downregulated genes, and **Supplementary Figure S2(A-Q)** provides the associated results.

**Figure 3.**
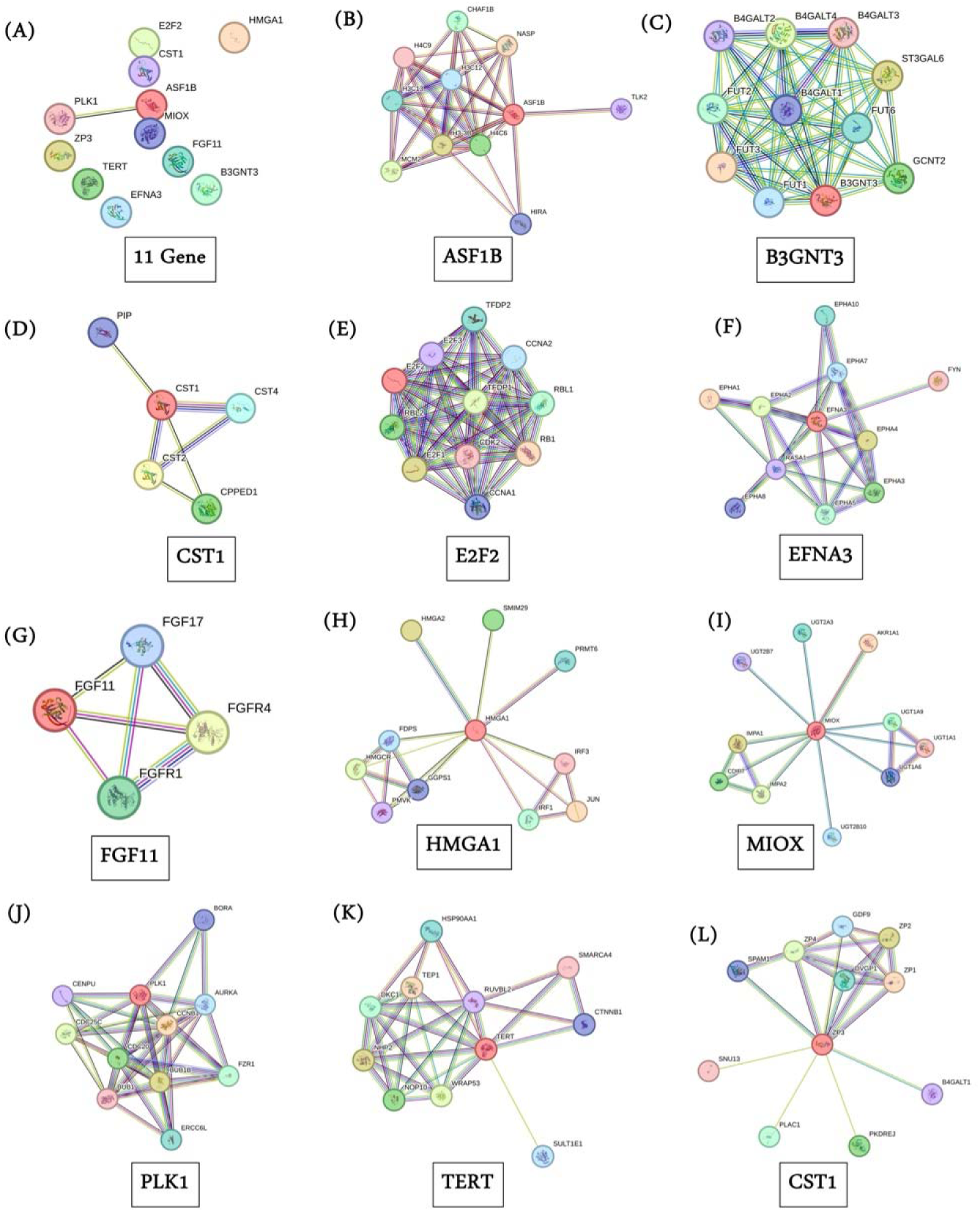
Protein-Protein Interaction (PPI) Analysis. Panel (A) represents PPI network generated using STRING database for the common 11 upregulated genes among MI, TCGA-LUAD and TCGA-LUSC datasets. Panel (B-L) represents PPI network generated using 11 genes individually.

### 3.4 Key genes are associated with overall survival in lung cancer patients

To investigate patients’ prognoses, we performed ‘Overall Survival’ analysis for the common up- and down-regulated genes using the GEPIA2 tool for both cancer types, LUAD and LUSC. Our analysis shows that all 11 common upregulated genes used as signatures were able to predict overall survival in LUAD in a statistically significant manner. The 11 gene signature show an HR of 1.8 with a log rank p-value of 8.3e-05 **[Fig. 4A]**. Similarly, our analysis of individual 11 genes revealed a significant association between 6/11 genes, namely *ASF1B, EFNA3, FGF11, HMGA1, PLK1,* and *ZP3*, and a poor prognosis in LUAD patients **[Fig 4B-G]** The average HR value observed was >1.4. The key role these genes play in tumorigenesis further explains this observation. For instance, Zhang et al. find a link between *ASF1B* and pathways related to cell proliferation in LAUD. They find that ASF1B indirectly controls the expression of CKS1B, POLE3, and DHFR, which ultimately leads to the progression of cancer [40]. Similarly, *FGF11* is associated with increased tumor proliferation in various cancer types, including LUAD [41], etc. A literature search suggested multiple pieces of evidence supporting our finding, in which they have shown how the above-mentioned genes are associated with poor prognosis in LUAD patients [41–44]. We found only *ASF1B* significantly associated with a poor prognosis in the case of LUSC **[Fig 4H]**. The Kaplan-Meir curve of the common upregulated genes with significant association is shown in **Fig. 4A–G** for LUAD and **Fig. 4H** for LUSC. KM curves for remaining upregulated common genes with non-significant association in predicting survival outcome are provided in **Supplementary Figure S3(A-E)** for LUAD and **Supplementary Figure S4(A-K)** for LUSC.

**Figure 4.**
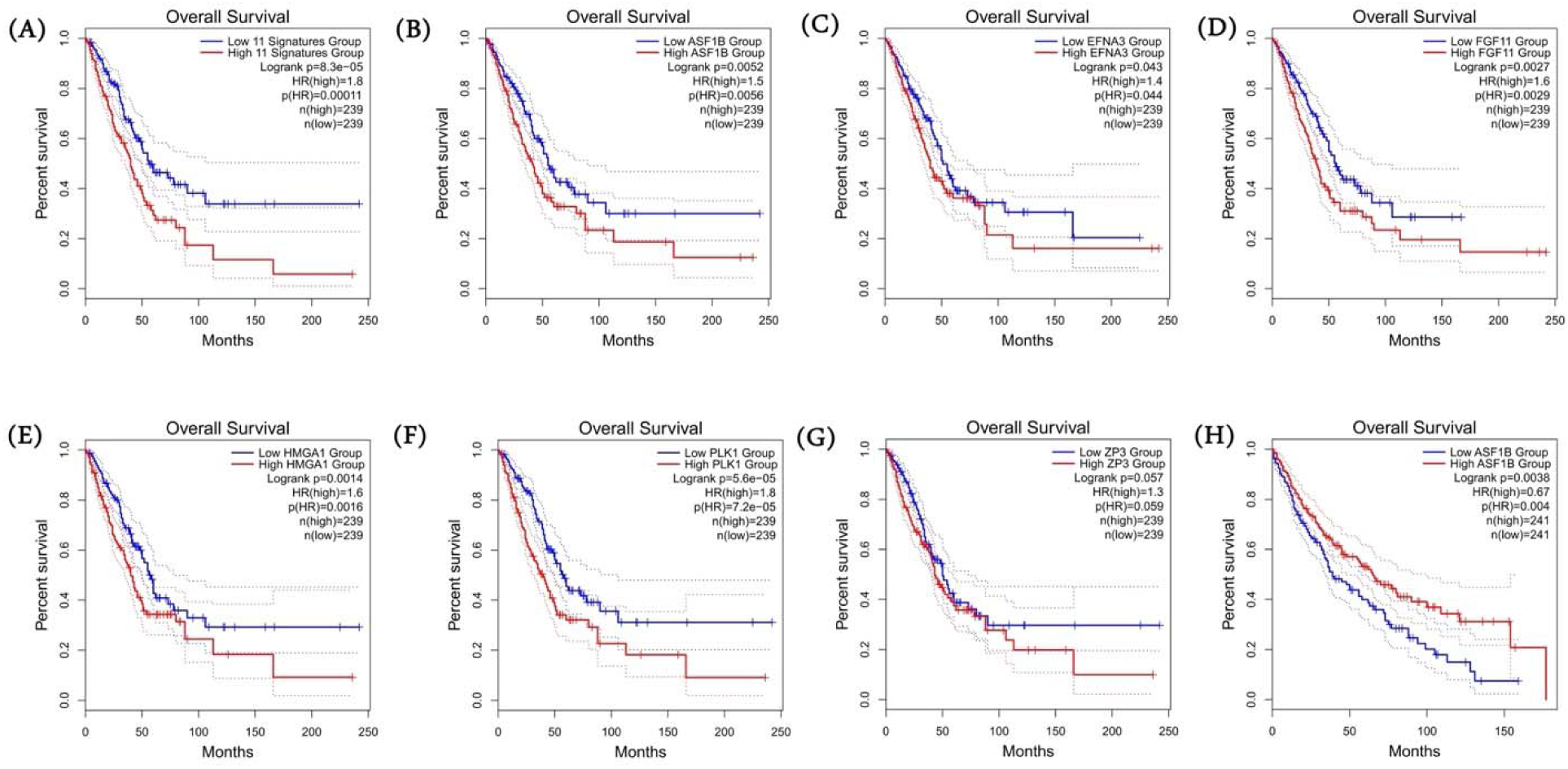
Common genes are associated with survival in lung cancer patients. Kaplan Meir (KM) Curve of the (A) 11 common upregulated genes among MI, TCGA-LUAD and TCGA-LUSC datasets as a ‘gene signature’ (B-G) individual genes which are statistically significantly as per log-rank test in TCGA-LUAD dataset. (H) KM curve of the gene ASF1B in TCGA-LUSC dataset. For each gene, patients were stratified into high and low category based on median expression.

In the case of common downregulated genes, all 16 genes, when used as signatures, didn’t significantly predict prognosis in LUAD. Upon examining each gene separately, we discovered that only three genes—*IL5RA, MS4A2, and SLC14A1*—were significantly associated with predicting overall survival in LUAD patients. These genes had a much lower hazard ratio (<1.0), indicating a better prognosis **[Fig. 5A–C]**. Previous studies revealed that these genes play a protective role in patients and validated our findings. Some researchers, like Fan et al. [45], found that *IL5RA* is linked to a better prognosis in LUAD patients. This might be because high levels of *IL5RA* are linked to an immune response against tumors. Likewise, Zheng et al. show that high expression of *MS4A2* genes is beneficial for the overall survival rate of LUAD patients [46]. *MS4A2* is associated with immune-infiltrating cells such as B cells, CD4^+^ T cells, CD8^+^ T cells, macrophages, etc., which play an important role in the immune response in patients. However, in LUSC, only *LRKK2* shows a significantly higher hazard ratio (>1.0), indicating a poor prognosis **[Fig. 5D]**. **Supplementary Figure S5 (A-N)** for LUAD and **Supplementary Figure S6 (A-P)** for LUSC provide KM curves for the remaining common downregulated genes, which show non-significant associations in predicting survival outcomes.

**Figure 5.**
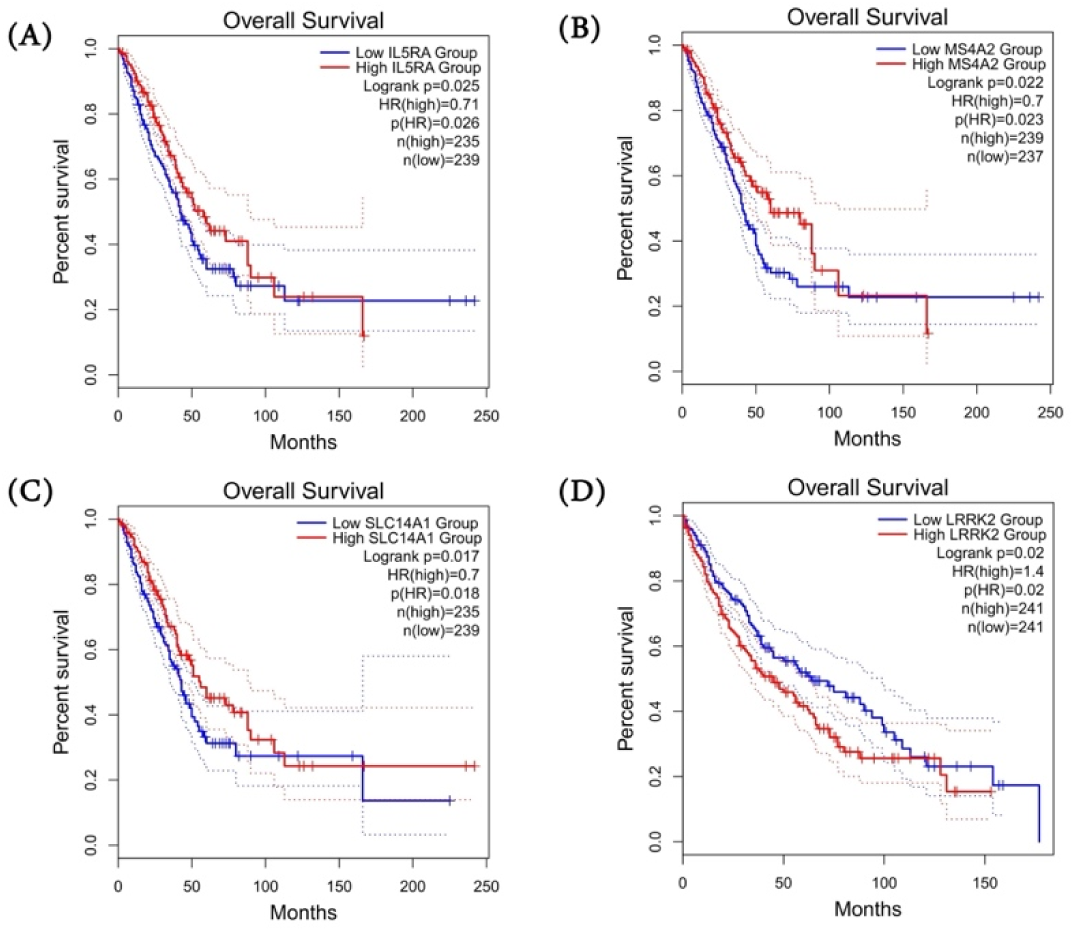
Kaplan Meir Curve analysis for commonly downregulated genes in lung cancer patients. Kaplan Meir (KM) Curve of the (A-C) individual genes which are statistically significantly as per log-rank test in LUAD dataset. (D) KM curve of the gene LRKK2 in LUSC dataset. For each gene, patients were stratified into high and low category based on median expression.

**Figure 6.**
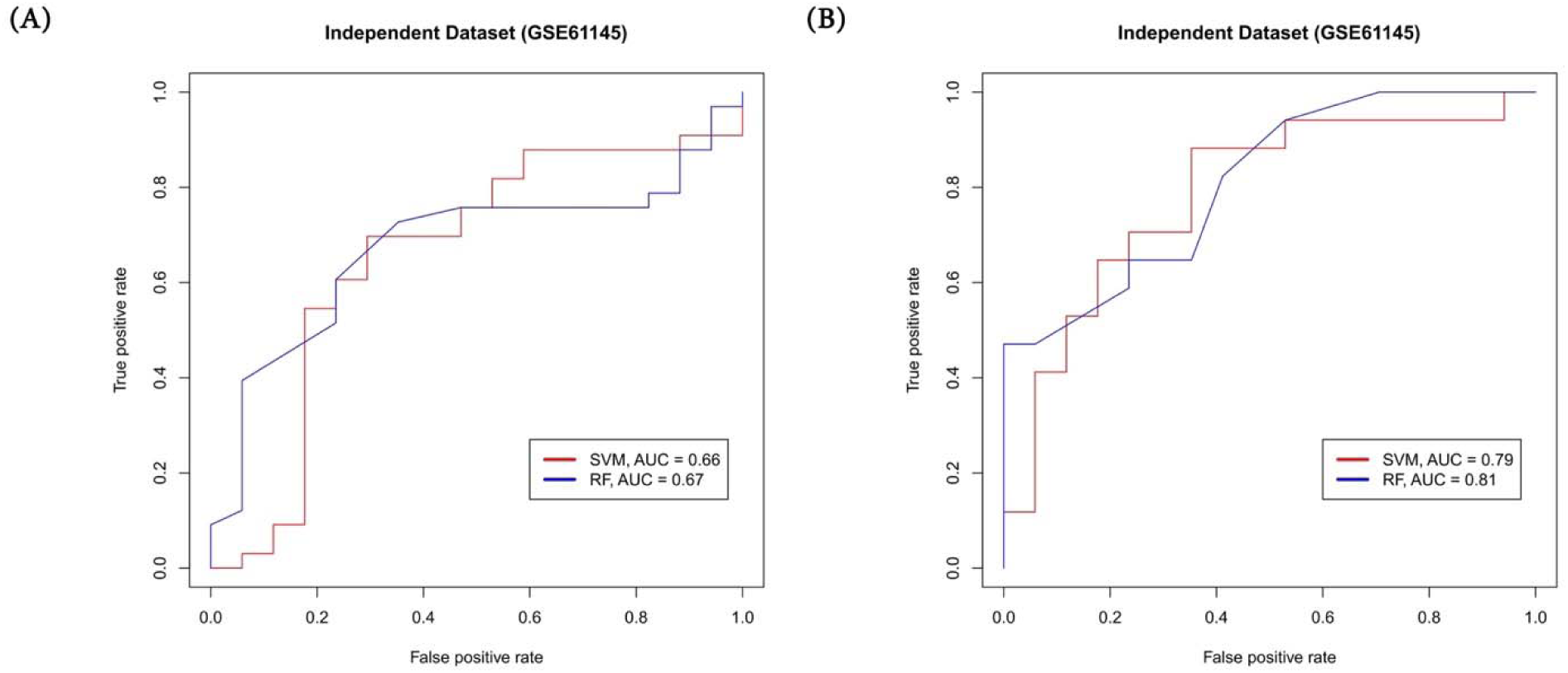
Performance of machine learning model on an Independent Dataset. Average gene expression of 24 common genes (11 up and 13 downregulated) obtained from 2 MI datasets, LUAD and LUSC were used as feature to build SVM and RF model to classify MI patients from normal. Model evaluation is shown on (A) Imbalanced dataset and (B) Balanced dataset.

### 3.5 Conserved site of miRNAs are present for the common genes among MI and lung cancer

TargetScanHuman 8.0 analysis, which used the common 11 upregulated genes, revealed one or multiple miRNAs with a conserved binding site for a given target gene. We observed conserved sites for six genes, which include *ASF1B, E2F2, EFNA3, FGF11, HMGA1,* and *TERT*. For the remaining 5 genes, there were no conserved sites for any miRNA. Next, we looked at the genes that had conserved sites. We discovered that E2F2 has 72 miRNAs with conserved sites, one more than any other gene. HMGA1 comes in second with 40 miRNAs, and EFNA3 comes in third with 28 miRNAs. We observed multiple miRNAs for each target and determined that the one with the lowest Context++ score and Seed Match Type (8mer, followed by 7mer-m8 and 7mer-1a) exhibited the strongest binding. **Table 3** lists the top 10 miRNAs for each target, selected based on the aforementioned criteria. For complete and detailed data, see **Supplementary Table S21**. We conducted a similar analysis for the 16 common downregulated genes, observing the conserved sites of miRNAs for only 10 targets. **Supplementary Table S22** provides the complete results. The overall analysis allows us to gain insight into miRNA-mRNA gene interaction and regulation, as well as design novel miRNA-based therapeutics.

**Table 3.**
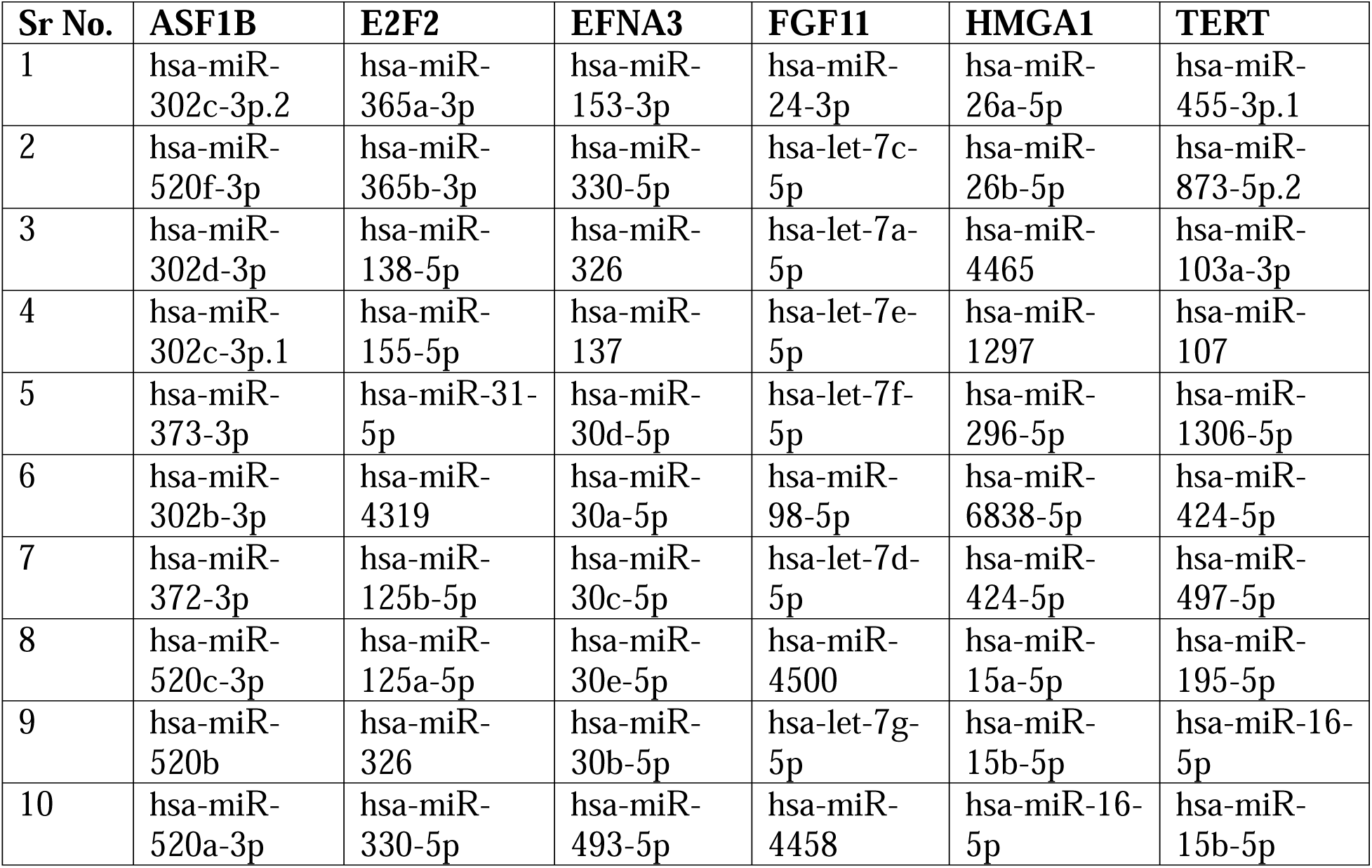
miRNA-mRNA relationship analysis. List of top 10 miRNAs having conserved site on mRNA characterized using TargetScanHuman 8.0.

### 3.6 Common genes can classify MI patients from normal with high accuracy

Firstly, we compared the gene expression of the common 11 upregulated and 16 downregulated genes in MI patients with their normal counterparts. As shown in **Supplementary Figure 7(A&B)**, we can clearly see a significant change in gene expression of 11 upregulated genes among MI patients compared to normal. Hence, we developed multiple machine learning models (SVM, and RF) that can classify MI patients from normal using the gene expression of the 27 common genes. We computed the average gene expression using the 2 MI datasets used in the study. We used a completely new and independent dataset, GSE61145 [47], which we did not use throughout this study, to validate the model performance. However, the validation dataset was missing data for the 3 genes (*ACKR4, FGFBP2,* and *HBA2*). Hence, we developed a 24-gene signature-based model and evaluated its performance on an independent dataset. We developed models using both imbalanced and balanced data. In the case of an imbalanced dataset, the RF classifier achieved the highest AUROC of 0.67, followed by SVM with AUROC of 0.66 **(Fig. 5A)**. However, on a balanced dataset, the RF-based model achieved the AUROC score of 0.81, followed by the SVM which had AUROC scores of 0.79 **(Fig. 5B)**. As there is no standard or previously reported gene signature associated with the discrimination of MI patients from normal, we couldn’t compare our model’s performance. We also looked at the gene expression of the 11 common upregulated genes in LUAD **[Supplementary Figure 8A]** and LUSC **[Supplementary Figure 8B]** and compared it to the normal in TCGA dataset. The Mann-Whitney Test showed that there was a significant difference in gene expression between the diseased population and the healthy population. Likewise, gene expression comparisons for the 16 downregulated genes in MI dataset is shown in **Supplementary Figure S9** and for LUAD and LUSC is shown in **Supplementary Figure S10**.

### 3.7 Drug Repurposing analysis potential drugs against MI and lung cancer

We conducted a drug repurposing analysis to identify potential drugs that target the 11 common upregulated genes associated with the disease biology in both MI and lung cancer. This was necessary because the current treatments have potential limitations, such as resistance and effectiveness, in a small population. Identifying new drugs that can target both MI and lung cancer patients could be beneficial for several reasons, such as shared molecular pathways, cost-effectiveness, comorbidity management, and so on. First, we map the target genes to UniProt, obtain the PDB ID, and download the 3D structure from the RCSB-PDB database. For genes lacking the PDB ID or full-length 3D structure, we download the AlphaFold predicted structure from the UniProt database. See **Supplementary Table S23** for all the details associated with the structures (PDB ID, chain, resolution, etc.). After downloading the structures, we used Open Babel software to process and refine them by removing heteroatoms and metal ions, among other things. Next, we put the refined structures on an online server called DrugRep (https://cao.labshare.cn/drugrep/). We then found the binding pockets on the protein surface and chose the one with the biggest volume because it has more active site residues that can interact during docking (**Supplementary Table S24)**. Next, we selected the ‘FDA-approved drugs’ library for screening against our target and submitted it for docking. After the docking experiment concluded, we downloaded a ‘tsv’ file containing the top 100 drugs, each represented by a ‘DrugBank ID’ and their free energy value. We mapped the top 3 hits with the highest free energy using the DrugBank database to obtain the drug names (**Supplementary Table S25**). **Table 4** provides a list of the top 3 drugs for each target.

**Table 4.** List of common 11 upregulated genes and top3 drugs mapped to them post drug repurposing analysis.

| Sr. No. | Target | Top 3 Drugs |
| --- | --- | --- |
| 1 | ASF1B | Lomitapide, Nebivolol, Conivaptan |
| 2 | B3GNT3 | Antrafenine, Telmisartan, Lomitapide |
| 3 | CST1 | Iloperidone, Sorafenib, Regorafenib |
| 4 | E2F2 | Conivaptan, Venetoclax, Ponatinib |
| 5 | EFNA3 | Dutasteride, Digitoxin, Ergotamine |
| 6 | FGF11 | Midosaurin, Bisotrizole, Adapalene |
| 7 | PLK1 | Tadalafil, Hexafluronium, Venetoclax |
| 8 | HMGA1 | Nilotinib, Conivaptan, Regorafenib |
| 9 | MIOX | Rupatadine, Trosipium, Rolapitant |
| 10 | TERT | Dihydroergocristine, Ergotamine, Dihydroergotamine |
| 11 | ZP3 | Darifenacin, Hydromorphone, Cyproheptadine |

## Discussion

Lung cancer is one of the leading malignant cancers in the world, affecting 2,480,675 people (both men and women) worldwide. As per the International Agency for Research on Cancer (IARC), lung cancer was the leading cause of cancer deaths, with 1.8 million deaths (18%) in 2020. Smoking is the leading cause of lung cancer, as it is responsible for nearly 85% of all cases [48]. One of the reasons for the high mortality rate in cancer is its diagnosis at an advanced stage, where treatment options are very few. If diagnosed at an early stage, the survival rate can be dramatically improved. Lung cancer has been hypothesized to be associated with increased cardiovascular diseases, especially coronary heart disease [49], stroke [50,51], and MI [20,52]. MI (also known as heart attack) is a type of CVD, that is prevalent worldwide and is associated with significant morbidity and mortality. MI occurs due to a decreased or complete cessation of blood flow to the myocardium. A large cohort-based study known as “INTERHEART” conducted across 52 countries identified several risk factors associated with MI, which include smoking, hypertension, abnormal lipid profile/blood lipoprotein (ApoB/ApoA1), diabetes mellitus, etc. [53,54].

Numerous studies have observed a direct or indirect association between lung cancer and MI [16,52,55,56]. They share similar risk factors, such as obesity, inflammation, and so on; however, because of their complex and distinct pathophysiology’s, analyzing both conditions has been challenging. In this study, we tried to address this issue by analyzing MI and lung cancer together. We performed differential gene expression analysis using genome-wide transcriptome data from GEO for MI patients and TCGA for lung cancer (LUAD and LUSC). We characterized DEGs (11 up- and 16 down-regulated) as common in all three conditions and performed several analyses. The GO analysis reveals an enrichment of pathways and processes found to be common among both diseased conditions. For example, common-up genes were found to be enriched for metabolic processes, immune-related processes, cell death regulation, IL-5-mediated signaling pathways, and so on, indicating a possible alteration in the metabolic and immune landscape in the tumor microenvironment and the cardiovascular system. Similarly, PPI interaction analysis using STRING reveals the association of common genes with other proteins that regulate a diverse range of interactions. Next, we looked for the survival association of the genes to predict prognosis in LUAD and LUSC patients and observed a significant association of the genes, especially Up genes in the LUAD cohort. The results demonstrate the prognostic value of the genes in cancer progression and open new avenues for developing targeted therapies. We also used common 11-up gene-based SVM and RF predictors to separate MI patients from the healthy population. We were able to get an AUROC of up to 0.99, which shows that these genes could be used as biomarkers for early diagnosis and risk assessment. Lastly, we performed drug repurposing analysis against the common 11-up genes and proposed novel drugs.

Up regulation of 11 genes that are common to both lung cancer and MI suggests a converging pathogenesis between lung cancer and cardiovascular events. This forms a rationale for the future work in the prediction of CV risk in patients with lung cancer, taking into account race and age-specific populations which will increase the predictive value of the computational platform with capabilities to predict patterns and processes associated with not only disease susceptibility, but also morbidity and mortality. Age and race are the shared risk factors in both lung cancer and MI cases due to the specific genetic predispositions and environmental exposures which can significantly increase the risk of disease in specific demographic groups [16–18]. Furthermore, the SVM and RF-based machine learning models developed in our study illustrated high accuracy (AUROC-0.99 and 0.95) of the common 11 genes in classifying MI patients from the healthy individuals. Also, Mann-Whitney Test depicted significant gene expression between diseased and healthy population in case of LUAD and LUSC indicating the potential to revolutionize personalized medicine by integrating these genetic biomarkers with clinical practice thereby improving early detection and intervention in high-risk populations. This also underscores the importance of investigating these gene signatures in populations with varied ethnic and genetic backgrounds, especially in Indian population, where the prevalence of tobacco consumption is high and influenced by varied socio-cultural practices [57]. The insights extracted from these molecular signatures regarding drug repurposing would indicate that there is a potential to develop targeted therapies that are based on the shared biological pathways of both lung cancer and MI. It is possible by taking into account genetic and demographic profiles of the patient population which will enable and enoble the foundation for precision medicine and precision public health in the field of CVD and Lung cancer. On a broader scale, the validation of these biomarkers in the race and age specific Indian population cohort would result in a novel screening tool that is more demographically and genetically pertinent to diverse communities, addressing health inequities in cardiovascular and cancer care.

Given that the current study suggests a potential link between MI and lung cancer pathophysiology, it’s important to address certain limitations. Firstly, we conducted the current study on a smaller dataset with limited geographical diversity; therefore, we must conduct the current analysis on a larger and more diverse cohort to confirm our findings. Second, we need to experimentally validate the proposed drugs after conducting a drug repurposing analysis. Third, we need to collect data where the patient with lung cancer has a previous history of MI or vice versa. Overall, the current study directs us to how we can leverage the commonality of the two diseases to improve our disease understanding and management, ultimately improving patient outcomes.

## Conclusion

The identification of significant upregulated genes from myocardial infarction (MI) and lung cancer, specifically LUAD, and LUSC, have suggested a possibility of potential shared mechanism of pathophysiology. Such genes could serve as potential biomarkers in early detection and risk stratification in both diseases, thereby improving triaging of patients and clinical decision-making. The application of machine learning models further supports the potential of these genetic signatures to classify patients with high accuracy, providing a promising tool for not only developing a sero-surveillance for LUAD/LUSC and MI patients in the community but also integrating precision medicine which will provide a base for precision public health as well. This paves the way to validate such biomarkers in larger and more diverse populations in a race specific manner for further assessment of the broader applicability of biomarkers, pursuing an ultimate goal to improve early detection, personalized treatments, and improved patient outcomes for both cardiovascular and lung cancer care. Also, our work provides a robust underpinning to further development in predicting the bi-directional risk of MI and lung cancer patients incorporating race- and age-specific variables. Demographic tailoring of such predictions will enhance the accuracy of algorithms using computational models that not only identify susceptibility to disease but also morbidity and mortality risks.

## Supporting information

All Supplementary Figures

All Supplementary Tables

## Acknowledgement

This research was supported by SRM Institute of Science & Technology, Kattankulathur, India.

## CRediT authorship contribution statement

**Dhruva Nandi:** Data Curation; Methodology; Writing Original Draft. **Rajiv Janardhanan:** Conceptualization and Writing Review & Editing. **Piyush Agrawal:** Conceptualization; Investigation; Data Curation; Methodology; Software; Formal analysis; Writing Original Draft; Writing Review & Editing; Project administration; and Supervision.

## Author contributions

PA collected and processed the dataset. PA, and DN performed the analysis. PA, and DN created the tables and figures. DN, RJ and PA wrote the manuscript. PA conceived the idea and supervised the study.

## Data and Code Availability

This paper analyzes existing, publicly available data. The accession numbers for the datasets used are listed in the key resources table. All the codes, datasets used to develop the machine learning models for each class is provided at out GitHub repository accessible at https://github.com/agrawalpiyush-srm/MI_LungCancer. Further information can be provided upon reasonable request to the corresponding author.

## Declaration of generative AI in scientific writing

During the preparation of this work the author do not used any AI or AI-assisted technologies to improve language and readability.

