## Supplementary material for "Predicting Prognostic Bidirectional Molecular Signatures Associating Myocardial Infarction and Lung Cancer: An *In-Silico* Perspective": All Supplementary Figures

Piyush Agrawal, Ph.D.

Division of Medical Research, SRM Medical College Hospital & Research Centre, SRMIST, Kattankulathur, Chennai, India-603203

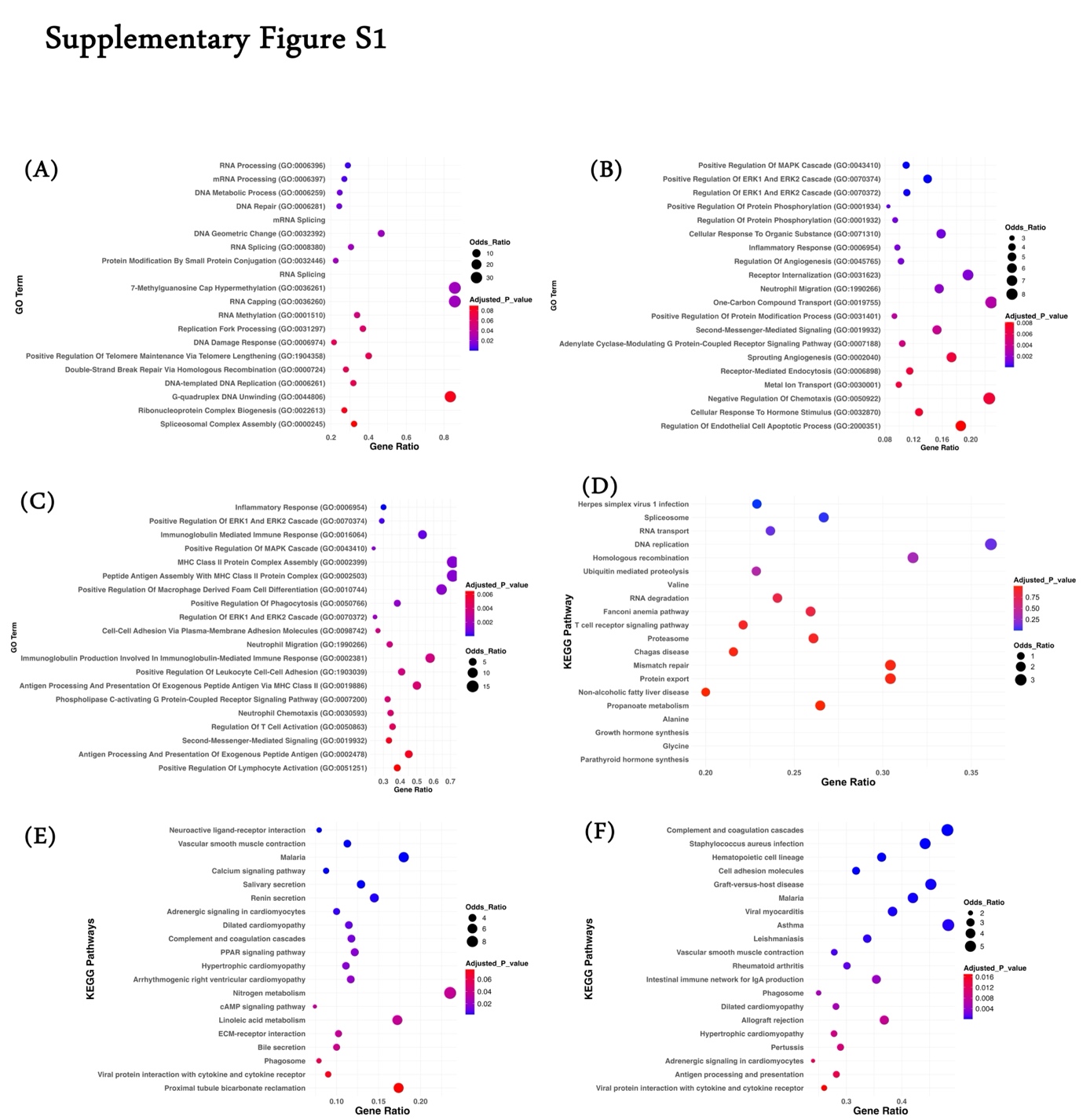

**Supplementary Figure S1.** **Gene Enrichment Analysis.** Panel A-C represents top20 statistically significant enriched biological processes associated with (A) Downregulated genes in MI datasets (combined); (B) Downregulated genes in TCGA-LUAD; (C) Downregulated genes in TCGA-LUSC. Panel D-F represents top20 statistically significant enriched KEGG pathways associated with (D) Downregulated genes in MI datasets (combined); (E) Downregulated genes in TCGA-LUAD; (F) Downregulated genes in TCGA-LUSC.

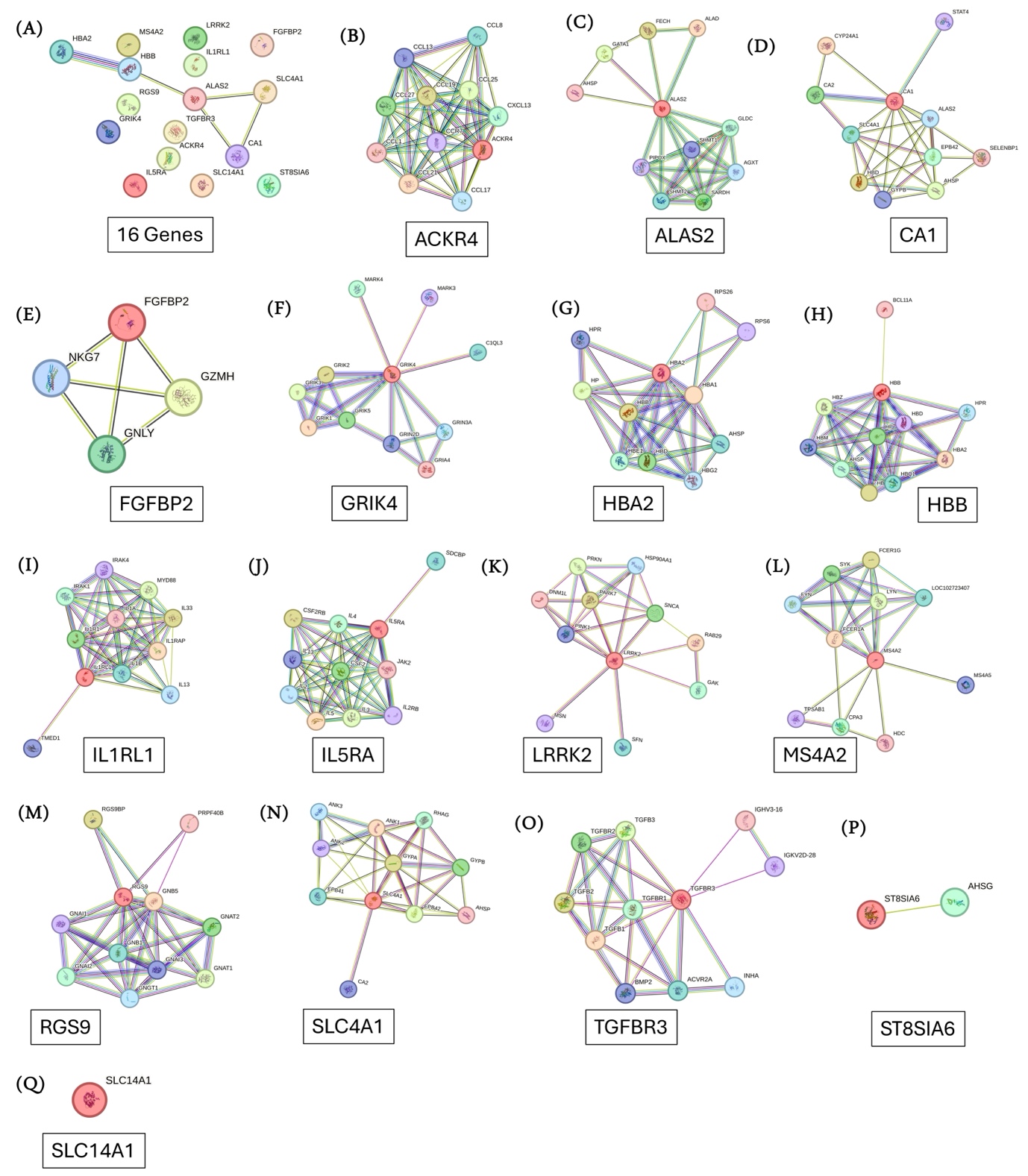

**Supplementary Figure S2. Protein-Protein Interaction (PPI) Analysis.** Panel (A) represents PPI network generated using STRING database for the common 16 downregulated genes among MI, TCGA-LUAD and TCGA-LUSC datasets. Panel (B-Q) represents PPI network generated using 16 genes individually.

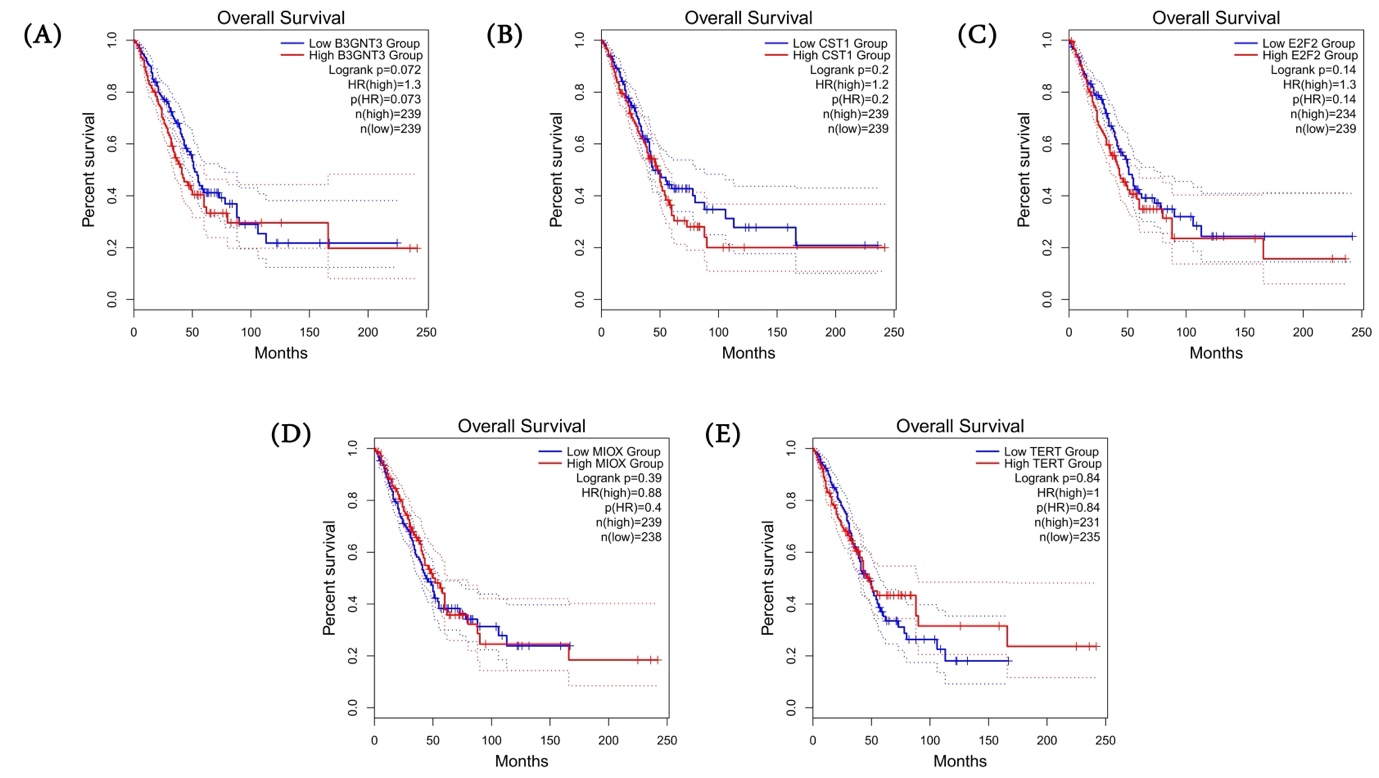

**Supplementary Figure S3. Kaplan Meir Curve Analysis.** Kaplan Meir (KM) Curve of the 5 common upregulated genes among MI, TCGA-LUAD and TCGA-LUSC datasets which are statistically insignificantly as per log-rank test in TCGA-LUAD dataset. For each gene, patients were stratified into high and low category based on median expression.

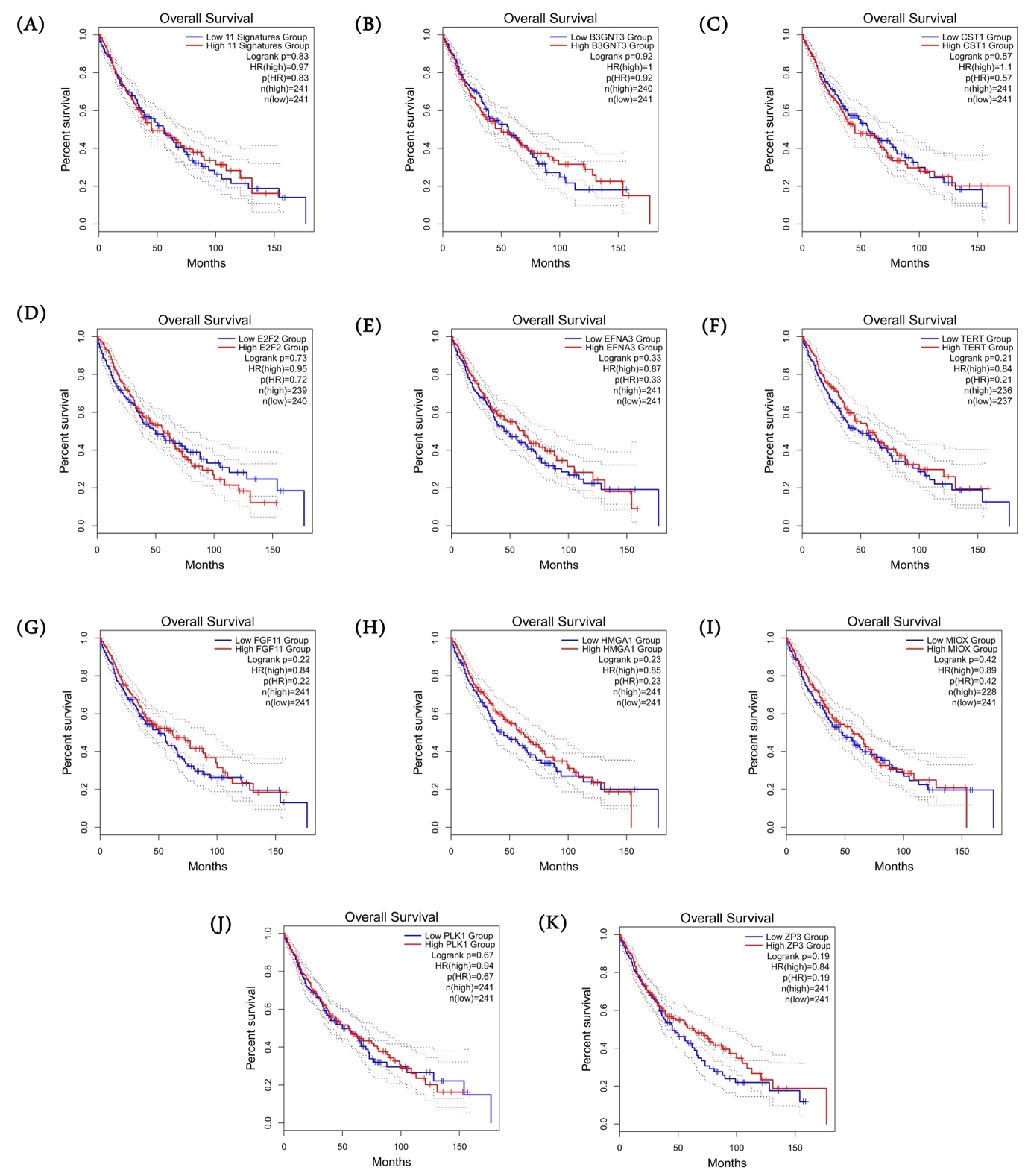

**Supplementary Figure S4. Kaplan Meir Curve Analysis.** Kaplan Meir (KM) Curve of the (A) 11 common upregulated genes among MI, TCGA-LUAD and TCGA-LUSC datasets as gene signature and (B-K) 10 individual genes which are statistically insignificantly as per log-rank test in TCAG-LUSC dataset. For each gene, patients were stratified into high and low category based on median expression.

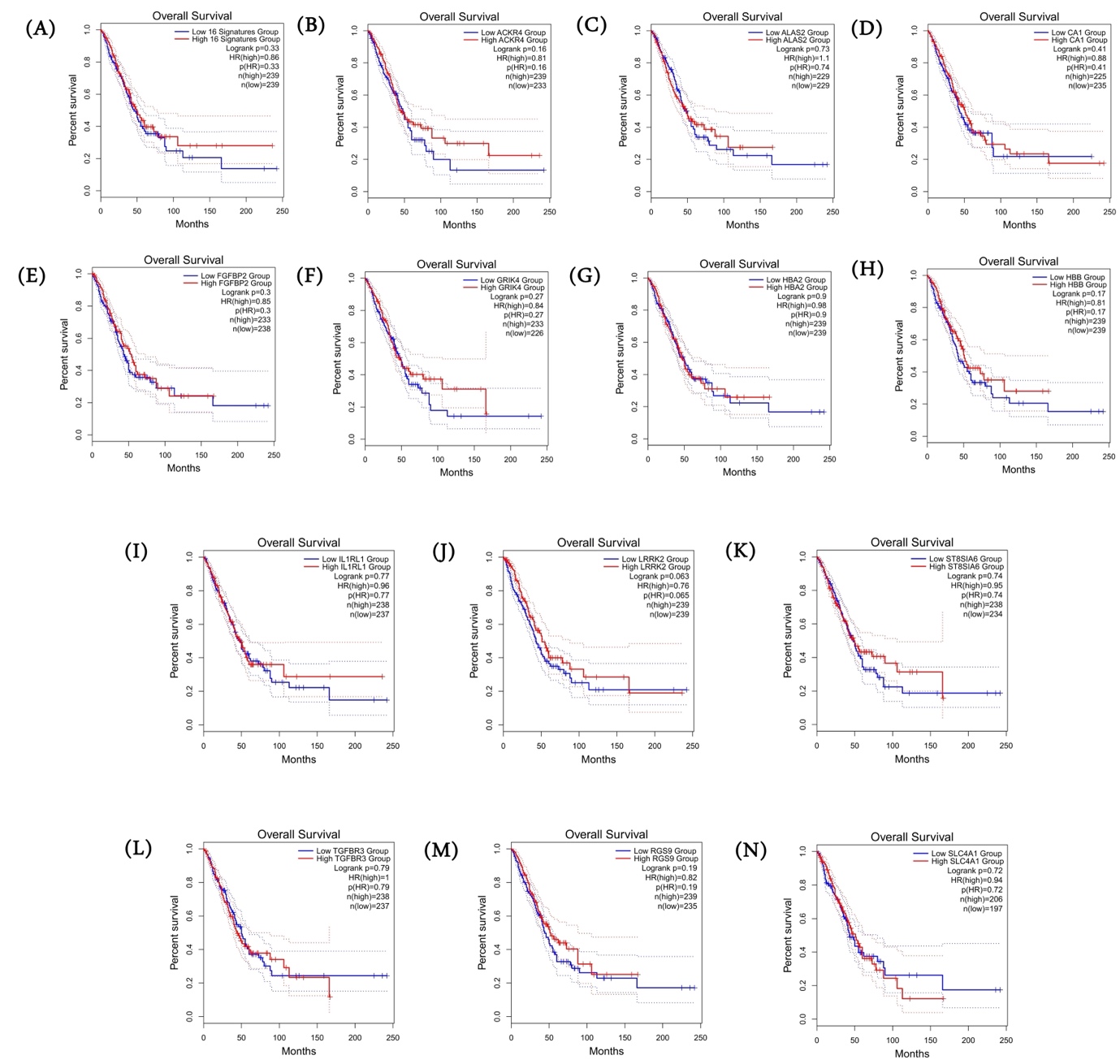

**Supplementary Figure S5. Kaplan Meir Curve Analysis.** Kaplan Meir (KM) Curve of the (A) 16 common downregulated genes among MI, TCGA-LUAD and TCGA-LUSC datasets as gene signature and (B-N) 13 individual genes which are statistically insignificantly as per log-rank test in TCAG-LUAD dataset. For each gene, patients were stratified into high and low category based on median expression.

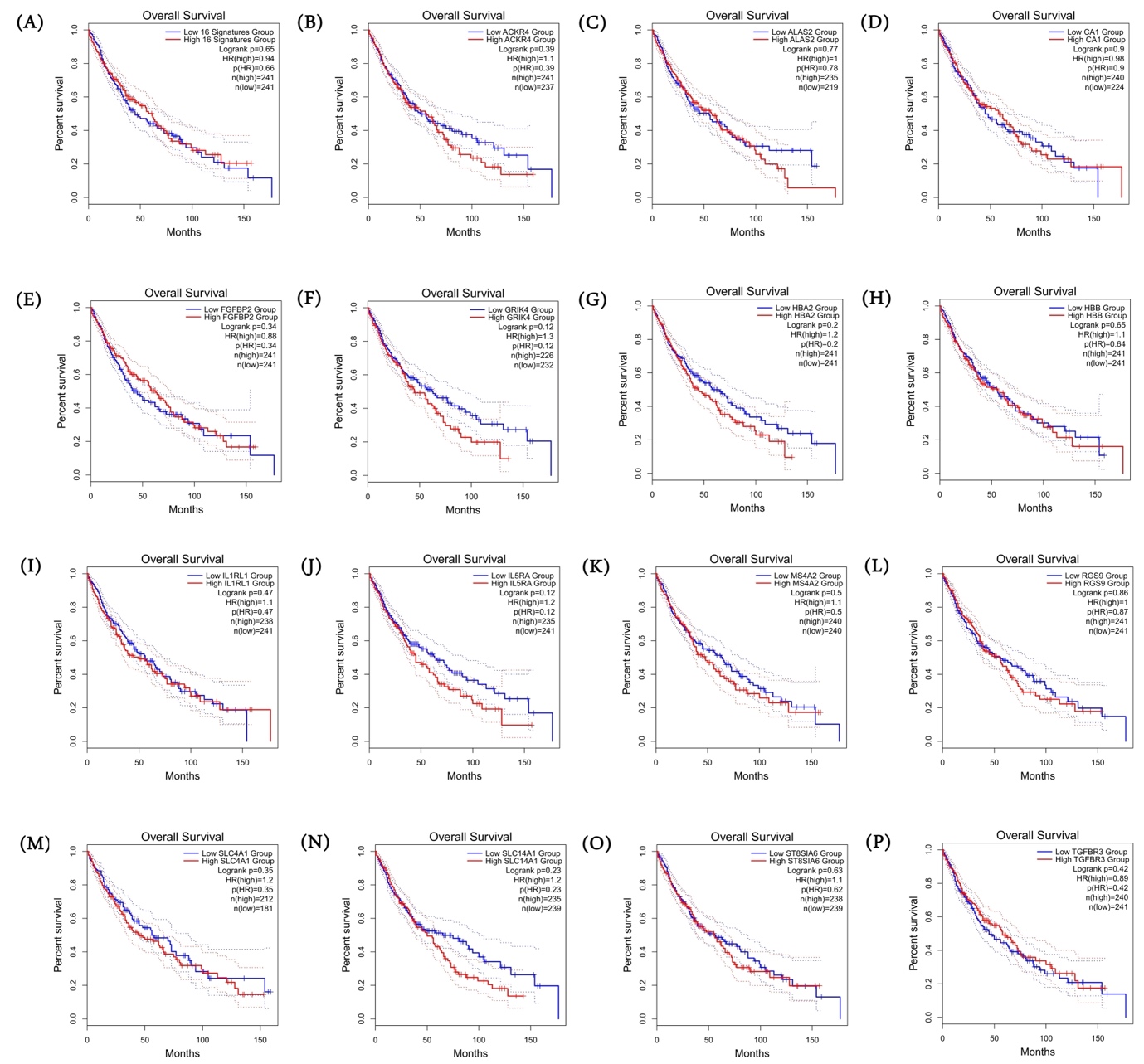

**Supplementary Figure S6. Kaplan Meir Curve Analysis.** Kaplan Meir (KM) Curve of the (A) 16 common downregulated genes among MI, TCGA-LUAD and TCGA-LUSC datasets as gene signature and (B-P) 15 individual genes which are statistically insignificantly as per log-rank test in TCAG-LUSC dataset. For each gene, patients were stratified into high and low category based on median expression

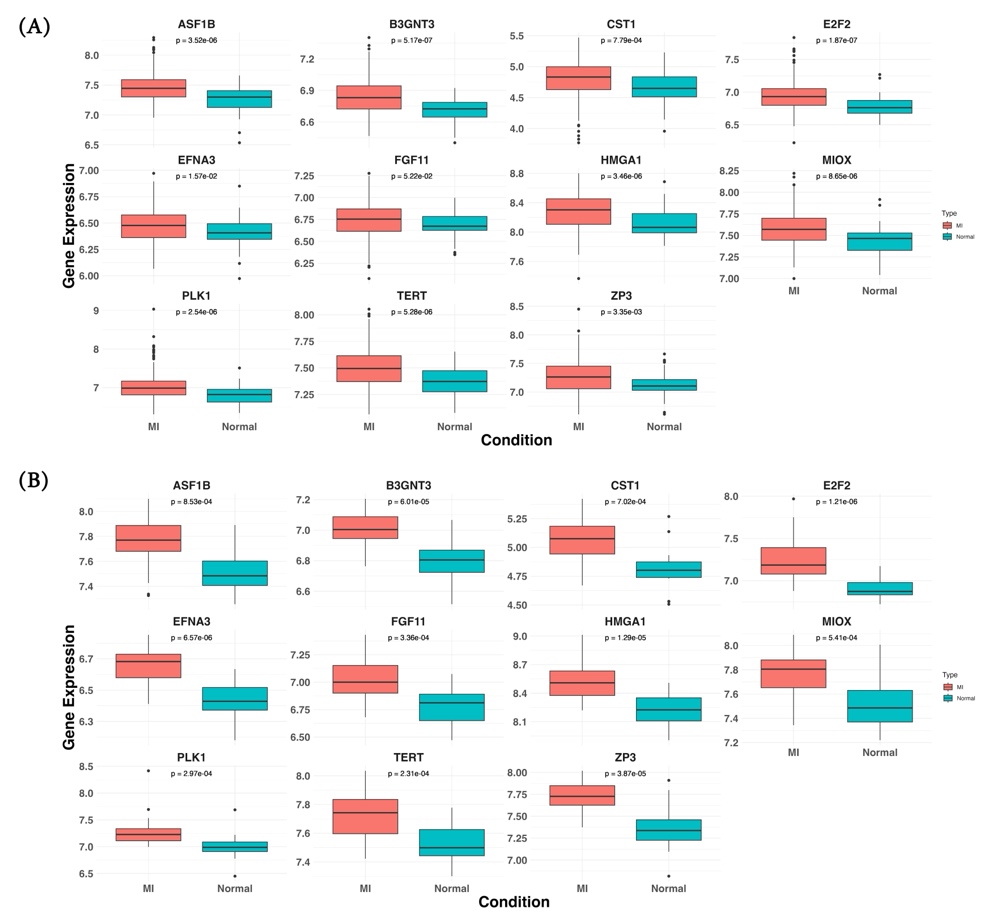

**Supplementary Figure S7. Gene Expression Comparison.** Gene expression comparison of the 11 common upregulated genes in MI patients with respect to control in (A) GSE59867 dataset and (B) GSE62646 dataset. Wilcoxon-test was performed to check the statistical significance.

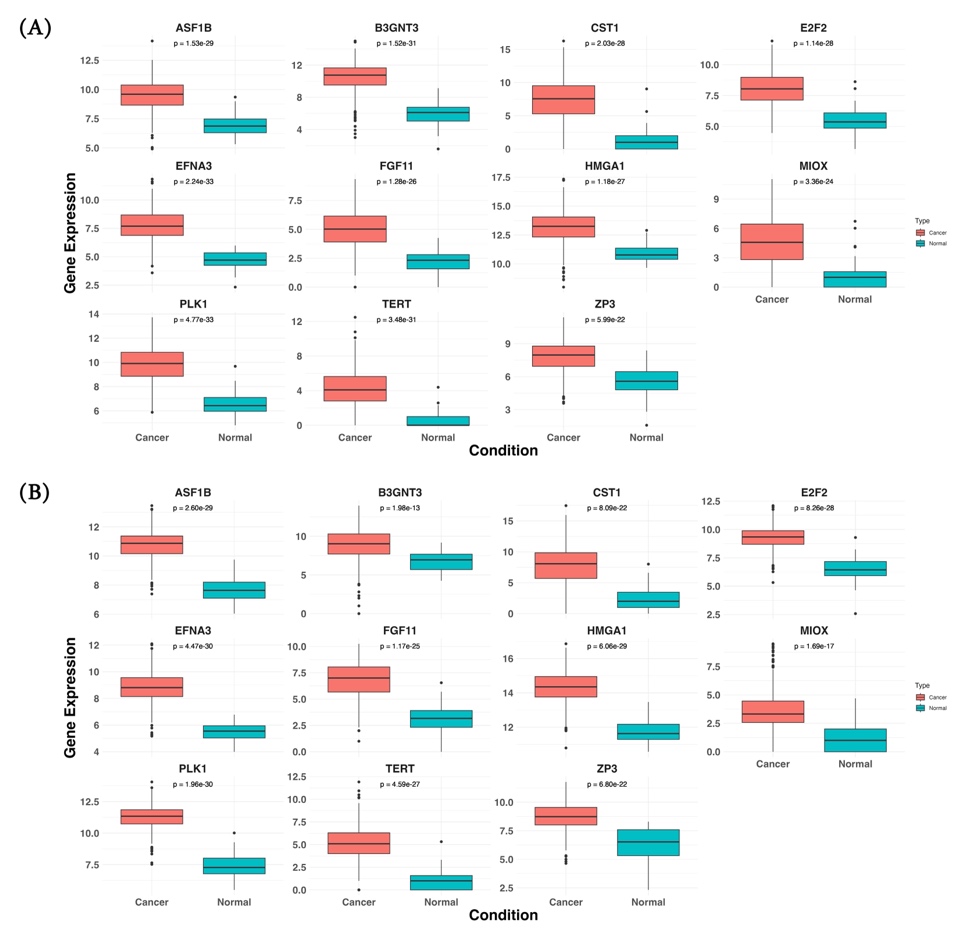

**Supplementary Figure S8. Gene Expression Comparison.** Gene expression comparison of the 11 common upregulated genes in (A) TCAG-LUAD dataset and (B) TCGA-LUSC dataset. Wilcoxon-test was performed to check the statistical significance.

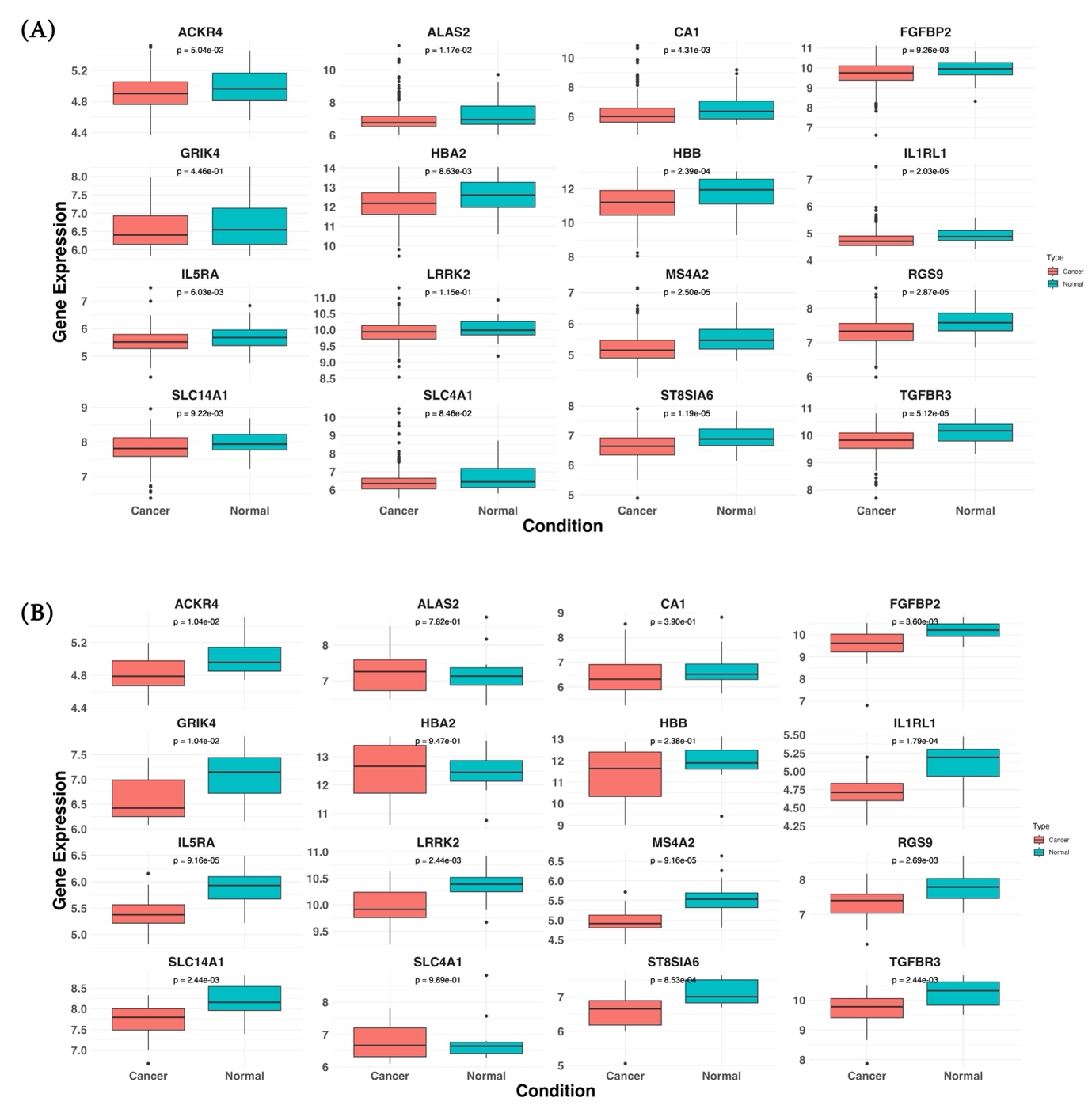

**Supplementary Figure S9. Gene Expression Comparison.** Gene expression comparison of the 16 common downregulated genes in MI patients with respect to control in (A) GSE59867 dataset and (B) GSE62646 dataset. Wilcoxon-test was performed to check the statistical significance

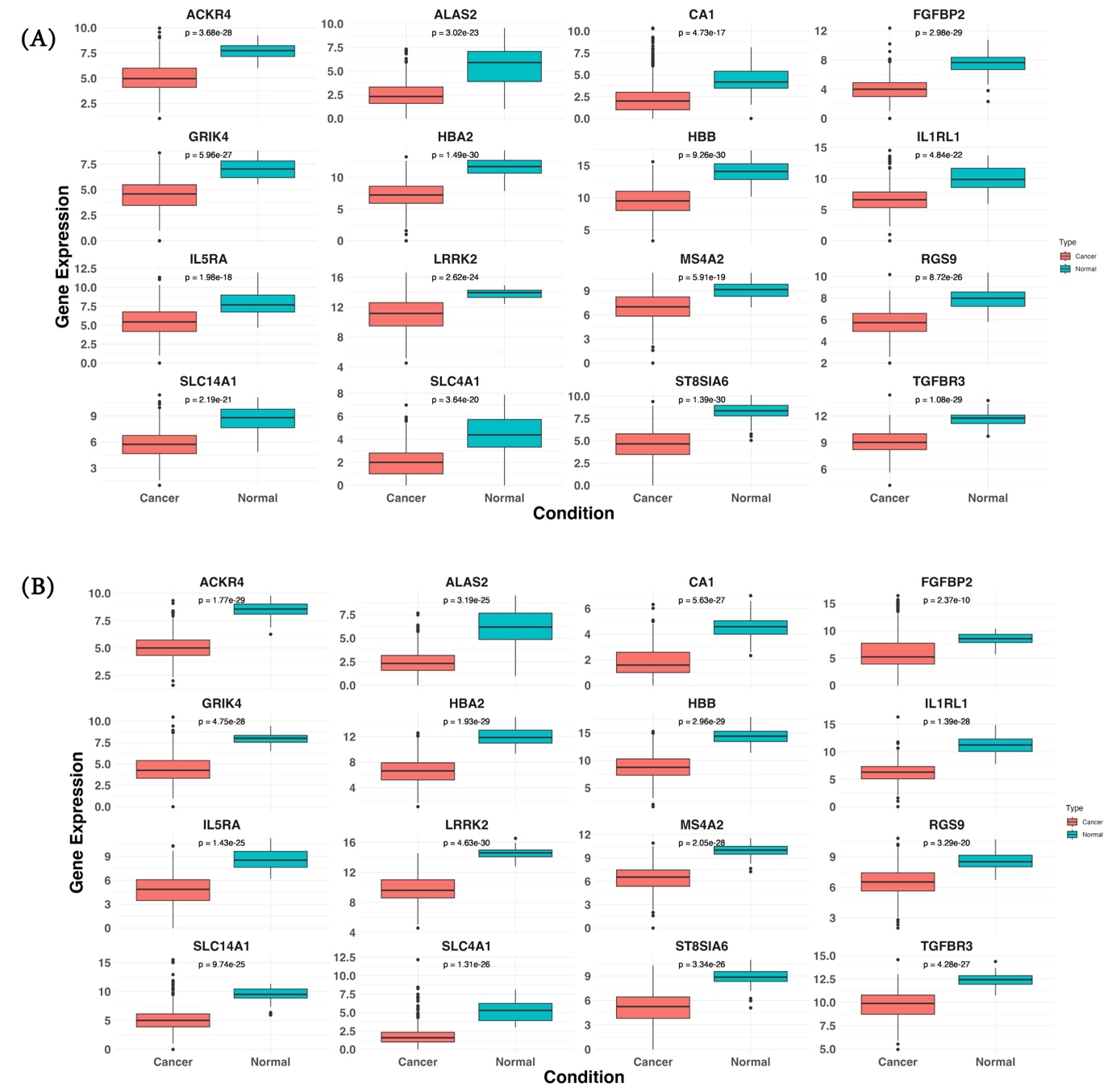

**Supplementary Figure S10. Gene Expression Comparison.** Gene expression comparison of the 16 common downregulated genes in MI patients with respect to control in (A) TCGA-LUAD and (B) TCGA-LUSC dataset. Wilcoxon-test was performed to check the statistical significance.
